# The nucleus measures shape deformation for cellular proprioception and regulates adaptive morphodynamics

**DOI:** 10.1101/865949

**Authors:** Valeria Venturini, Fabio Pezzano, Frederic Català Castro, Hanna-Maria Häkkinen, Senda Jiménez-Delgado, Mariona Colomer-Rosell, Mónica Marro Sánchez, Queralt Tolosa-Ramon, Sonia Paz-López, Miguel A. Valverde, Pablo Loza-Alvarez, Michael Krieg, Stefan Wieser, Verena Ruprecht

## Abstract

The physical microenvironment regulates cell behavior during tissue development and homeostasis. How single cells decode information about their geometrical shape under mechanical stress and physical space constraints within their local environment remains largely unknown. Here we show that the nucleus, the biggest cellular organelle, functions as a non-dissipative cellular shape deformation gauge that enables cells to continuously measure shape variations on the time scale of seconds. Inner nuclear membrane unfolding together with the relative spatial intracellular positioning of the nucleus provides physical information on the amplitude and type of cellular shape deformation. This adaptively activates a calcium-dependent mechano-transduction pathway, controlling the level of actomyosin contractility and migration plasticity. Our data support that the nucleus establishes a functional module for cellular proprioception that enables cells to sense shape variations for adapting cellular behaviour to their microenvironment.

**One Sentence Summary:** The nucleus functions as an active deformation sensor that enables cells to adapt their behavior to the tissue microenvironment.

## Main Text

The 3D shape of an organism is built by active force-generating processes at the cellular level and the spatio-temporal coordination of morphodynamic cell behavior. Contractility of the acto-myosin cell cortex represents a major cellular force production mechanism underlying cellular shape change (1), cell polarization (2) and active cell migration dynamics (3). Contractility levels are regulated by the activity of non-muscle myosin II motor proteins (4) and are spatio-temporally controlled to tune single cell and tissue morphodynamics during development (5, 6) and tissue homeostasis and disease in the adult organism (7, 8). Still, mechanisms that regulate the set point level of cortical contractility on the single cell level remain poorly understood.

To adjust cortical contractility levels, cells need to make quantitative measures of their mechano-chemical 3D tissue microenvironment and translate this information into a defined morphodynamic output response. During embryogenesis, morphogens that act as chemical information carriers have attracted major attention (9), modulating cytoskeletal and cellular dynamics via receptor signaling pathways that tune protein activities (such as phosphorylation states) and/or protein expression levels. In contrast, physical parameters of the 3D tissue niche and mechanical forces gain importance as regulators of cellular morphodynamics and myosin II-dependent cortical contractility levels (10, 11). In vivo, mechanical cell deformation and cellular packing density in crowded tissue regions has been shown to influence major morphodynamic processes such as cortical actomyosin contractility (12, 13), cell division (14), cell extrusion (15, 16) and invasion (17). Ex vivo studies further provided evidence on the single-cell level that physical cell deformation is sufficient to modulate cortical myosin II localization and motor protein activity (18, 19) and influencing morphodynamic cell behavior (20, 21).

A recent example has been the identification that fluctuations in cortical contractility by genetic or physical cell perturbation are sufficient to induce spontaneous cell polarization in embryonic stem cells (22), underlying the induction of a fast amoeboid stable-bleb migration mode. This morphodynamic migration switch was shown to be present in both undifferentiated and lineage committed embryonic progenitor cells and was further identified in various differentiated cell types and transformed cancer cell lines (23, 24). This suggests that a conserved, yet unknown, mechanosensitive cellular signaling module regulates myosin II-based cortical contractility and cell transformation depending on cellular shape deformations in constrained tissue microenvironments.

To approach the question of how cells can measure and adaptively respond to physical cell shape changes within their 3D tissue microenvironments, we established a synthetic approach that enables to mimic mechanical cell deformations in controlled 3D microconfinement assays (25). Primary progenitor stem cells were isolated from blastula stage zebrafish embryos and cultured in planar confinement assays of defined height to mimic various cell deformation amplitudes (Fig. S1A). Lowering confinement height in discrete steps increased cell deformation, which scaled non-linearly with a pronounced enrichment of myosin II at the cell cortex relative to cortical actin accumulation (Fig.1A,B and Fig. S1B,C; Movie 1). Cortical accumulation of myosin II was accompanied by an increase in cellular bleb size (Fig. S1D, Movie 1), indicative of an active increase in cortical contractility levels depending on confinement height. Myosin II re-localization to the cell cortex in confined cells was rapid (t_1/2_ < 1min, Fig. 1C, D) and temporally stable under confinement. Of note, distinct plateaus of cortical myosin II enrichment were evident, with myosin II re-localization increasing for larger cell deformations (Fig. 1C). A cell confinement height below 7 μm caused a pronounced increase in cell lysis during compression, defining a maximal threshold deformation of ∼30% of the initial cell diameter, given a blastula cell size of d∼25 μm (Fig. S3G). Overall, these data support that the physical microenvironment defines a specific set point level of cortical contractility as a function of physical cell deformation in confined microenvironments.

**Fig. 1.**
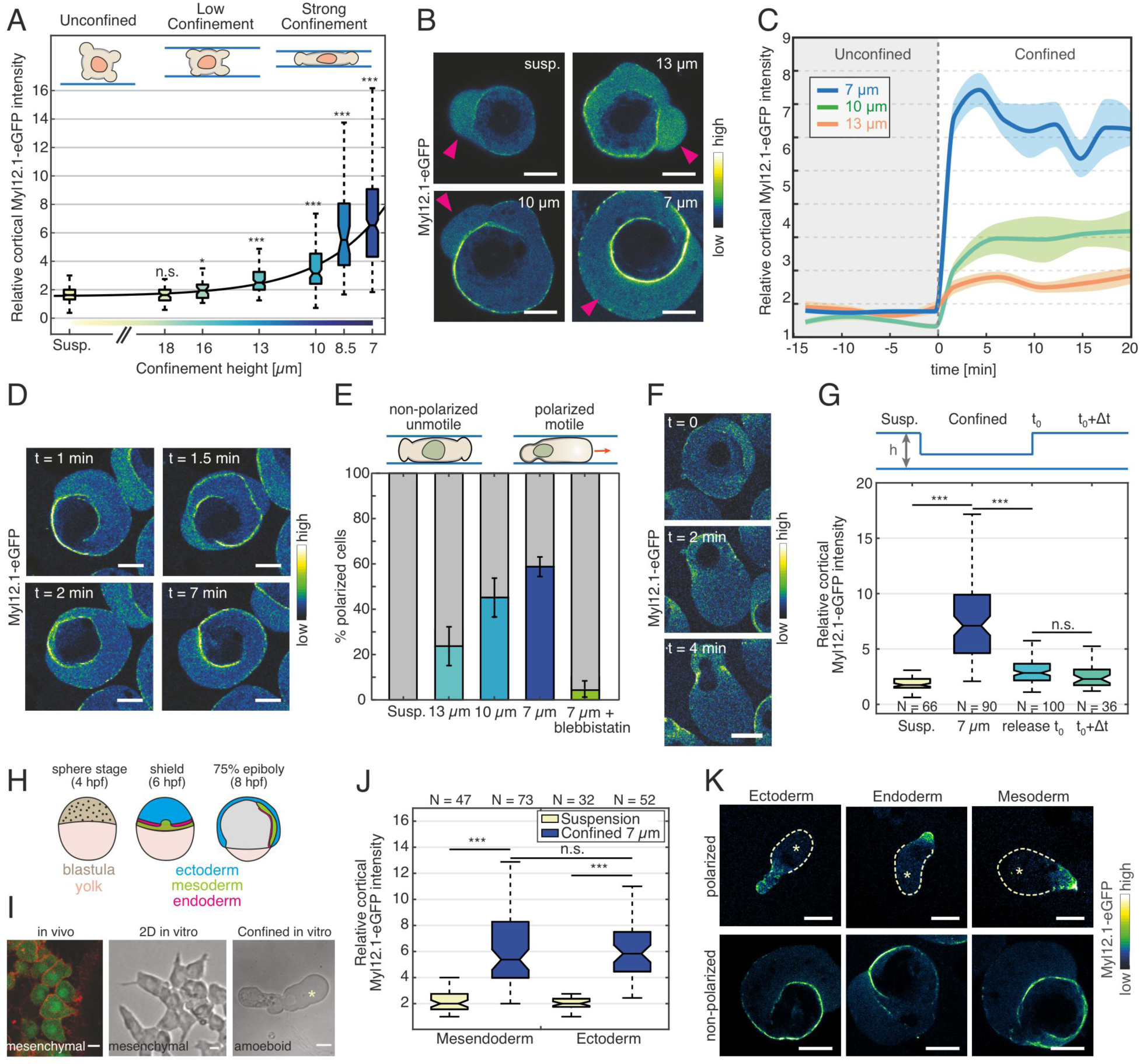
Cell deformation in confined environments defines cortical contractility, polarization and fast amoeboid cell migration. **(A)** Relative cortical myosin II enrichment for decreasing confinement height in un-polarized progenitor cells (n=477 (suspension, unconfined); n=56 (18 μm); n=35 (16 μm); n=103 (13 μm); n=131 (10 μm); n=49 (8.5 μm); n=348 (7 μm)). Significance values are with respect to the suspension condition. Black line shows a mono-exponential fit with offset to the data. **(B)** Exemplary confocal fluorescence images of control progenitor stem cells in suspension (Susp., unconfined) and confinement at indicated heights expressing Myl12.1-eGFP. Magenta arrows point at cellular blebs. **(C)** Temporal dynamics of cortical myosin II recruitment upon mechanical confinement at time t=0 at indicated heights. **(D)** Exemplary confocal fluorescence time-lapse images of cells expressing Myl12.1-eGFP under 7 μm confinement. **(E)** Percentage of polarized motile stable-bleb cells in suspension, indicated confinement heights and 7 μm confinement with 10 μM Blebbistatin (each n>500). **(F)** Exemplary time-lapse images of a cell expressing Myl12.1-eGFP undergoing spontaneous cell polarization. **(G)** Relative cortical myosin II enrichment during reversible cell confinement. Cells were confined for 15 min before confinement was released and cortical myosin II levels were measured at t_0_ (0-5 min) and at t_0_+Δt (30-60 min) after release. **(H)** Sketch of the developing zebrafish embryo at sphere (4 hpf), shield (6 hpf) and 75% epiboly (8 hpf) stage. **(I)** Exemplary confocal and bright field images of mesodermal cells in vivo expressing Lyn-Tomato (membrane) and GFP under the mezzo promotor (left), induced mesendodermal cell in vitro plated on a 2D fibronectin-coated surface (middle) and under 7 μm confinement (right). **(J)** Relative cortical myosin II intensity for induced mesendodermal and ectodermal progenitor cells in control suspension and confinement conditions. **(K)** Exemplary confocal images of stable-bleb polarized (top) and non-polarized (bottom) cells expressing Myl12.1-eGFP under 7 μm confinement. From left to right: induced ectoderm endoderm and mesoderm cells. Yellow asterisks indicate the front of polarized stable-bleb cells. ***p<0.001, **p<0.01, *p<0.05, not significant (n.s.). All scale bars 10 μm.

We have previously shown that an increase in myosin II-mediated cortical contractility induced a stochastic motility switch into a highly motile amoeboid migration phenotype termed stable-bleb mode (22). In accordance with these results, rapid cortical myosin II enrichment in confinement resulted in spontaneous cell polarization (Fig. 1E,F and Fig. S1J, Movie 2). Polarized cells revealed characteristic actomyosin density gradients from the cell front towards the rear (Fig. S1E), accompanied with fast retrograde cortical flows (Fig. S1F, Movie 3) that have been shown to power cell locomotion (22, 26) and induce fast amoeboid migration in polarized cells (Fig. S1H, I). These data show that physical cell deformation in confinement is sufficient to increase actomyosin network contractility and trigger rapid amoeboid cell migration, mimicking the morphodynamic phenotype observed under ectopic activation of myosin II. The fraction of polarized cells and associated migration rate strongly increased with cell confinement, suggesting that large cell deformations can promote cell polarization and active cell migration as a cellular response to shape changes in constrained tissue environments.

Release of cell compression in confinement induced a rapid re-localization of cortical myosin to the cytoplasm, followed by a rapid loss of cell polarization and associated migratory capacity (Fig. 1G and Fig. S1G, Movie 4). Interfering with myosin II activity via Blebbistatin inhibited cell polarization and motile transformation in confinement (Fig. 1E and Fig. S1J), in accordance with a necessary role of myosin II-based contractility in cell polarization and migration induced by mechanical cell shape deformation. Cortical myosin II enrichment and cell polarization occurred independently of caspase activation (Fig. S1K), supporting that morphodynamic changes are not caused by the activation of pro-apoptotic signaling programs.

During gastrulation, blastoderm embryonic progenitor stem cells specify into different lineages (ectoderm, mesoderm, endoderm) and acquire distinct biomechanical and morphodynamic characteristics, driving germ layer positioning and shape formation of the embryo (Fig. 1H). To test the mechanosensitive response to cell deformation at later developmental stages, we obtained different progenitor cell types from embryos via genetic induction or using endogenous reporter lines. Under confinement, non-motile ectodermal cells rapidly polarized and started to migrate, similarly to mesendodermal cells that underwent a fast mesenchymal-to-amoeboid transition in confinement (Fig. 1I-K and Fig. S2A,B, Movie 5), supporting that physical shape constraints are sufficient to induce rapid changes in cortical contractility and migration types, irrespective of cell fate specification and initial migratory programs. Together, these results support that physical cell shape deformation in confined tissue microenvironments activates a mechanosensitive signaling pathway regulating adaptive cortical contractility levels and morphodynamic migration plasticity in pluripotent and lineage committed embryonic stem cells.

We next sought to identify potential mechanisms that control cellular shape deformation sensing and adaptive morphodynamic behavior. Cortical myosin re-localization occurred on passivated confinement surfaces independently of adhesive substrate coating (Fig. S2B, 3A) and cell-cell contact formation (Fig. S3B). These observations support that the activation of cortical contractility in confinement occurs independently of adhesion-dependent mechano-transduction pathways (27). The temporal characteristics of myosin re-localization in confined cells (fast, stable and reversible accumulation of cortical myosin), suggest that shape deformation is sensed by a non-dissipative cellular element that can rapidly measure and convert gradual cellular shape changes into a stable contractility response levels.

The actomyosin cytoskeleton itself has been implicated to act as a mechanosensitive network (19), but is generally limited in sustained deformation sensing due to rapid turnover of the cell cortex (28). Testing for the activation of mechanosensitive ion channels using GsMTx4 (an inhibitor of the tension-dependent Piezo1 channel that is activated following confinement of human cancer cells (29)) showed no significant reduction in cortical myosin accumulation under cell deformation (Fig. S3C), despite the presence of functional Piezo1 channels in these cells (Fig. S3D). Interestingly, we observed that cortical myosin II enrichment only started to occur below a threshold confinement height (∼13 μm) that correlated with the spatial dimension of the nucleus (Fig. 2A and Fig. S3G). Analyzing nuclear shape change versus cortical myosin accumulation revealed a bi-phasic behavior, with a first phase in which the nucleus diameter remained nearly constant and no myosin accumulation was observed, and a second phase in which the relative myosin accumulation linearly increased with the relative change in nucleus diameter (Fig. 2A,B). In accordance with this observation, we expected a proportional change of nuclear surface ruffling upon deformation of an initially spherical nucleus. Measuring of nuclear surface folding by the expression of the inner nuclear membrane (INM) protein Lap2b-eGFP revealed that membrane ruffling was continuously reduced when nucleus deformation started to occur at a threshold deformation of ∼13 μm (Fig. 2C-E, Fig. S3E, Movie 6). In addition, analysis of nucleus membrane curvature for confined versus control cells in suspension indicated INM surface unfolding that remained stable over the measurement time of 60 min (Fig. 2F,G, Movie 6), with no significant difference in total nuclear volume and surface (Fig. S3F). Nucleus deformation further correlated with cortical myosin II accumulation in the endogenous in vivo context during the blastula to gastrula transition, when a gradient of cellular packing density appears from the animal pole towards the lateral margin (30) (Fig. S2C,D).

**Fig. 2.**
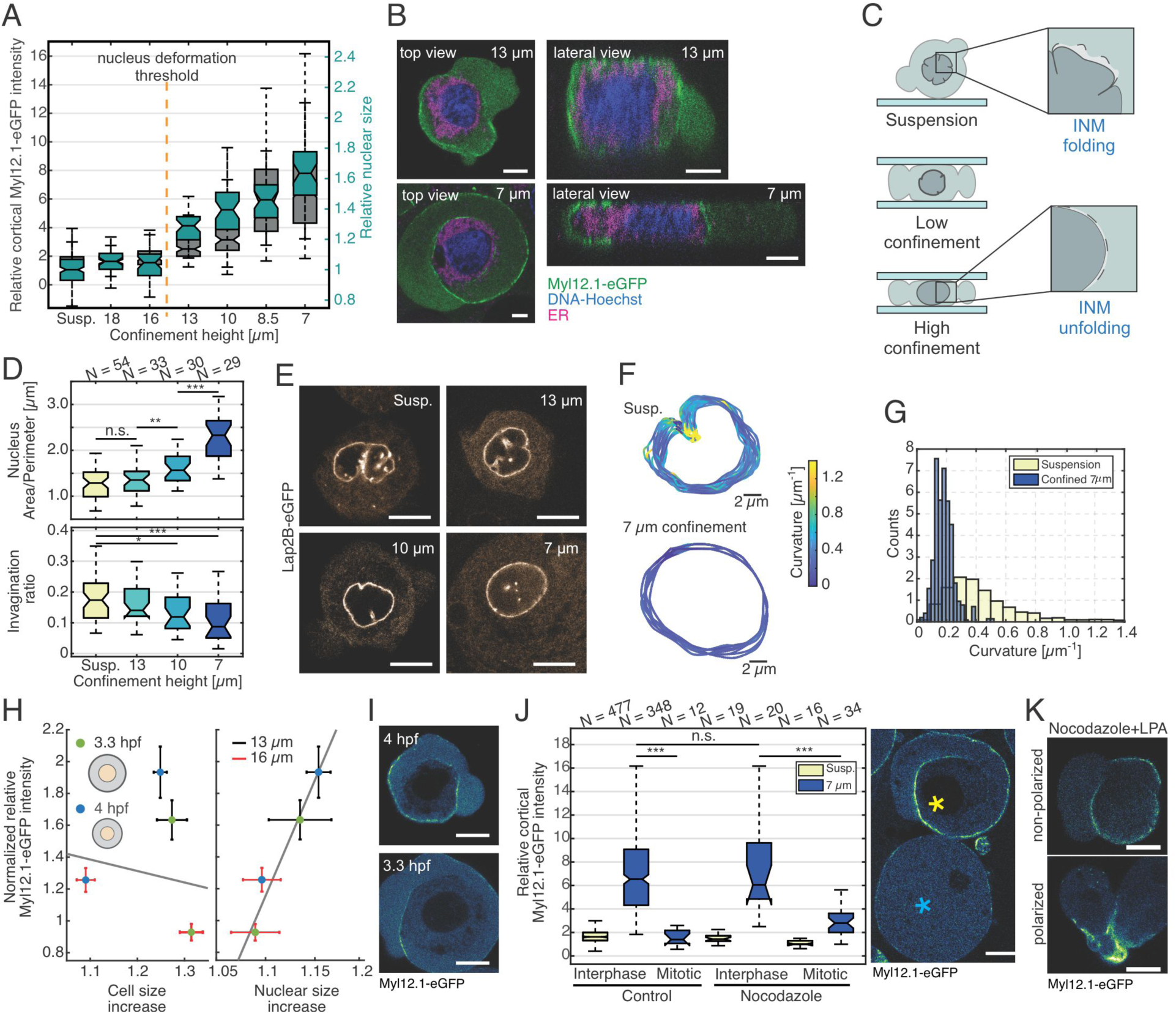
Nuclear envelop unfolding is associated with increasing cortical contractility. **(A)** Double boxplot of relative cortical myosin II enrichment (left axis, green) and nuclear size increase (right axis, grey) for decreasing confinement height. **(B)** Exemplary confocal top views (x-y) and side views (y-z) of progenitor stem cells expressing Myl12.1-eGFP stained with DNA-Hoechst and ER-TrackerRed for 13 μm and 7 μm confinement. **(C)** Illustration showing the unfolding of the inner nuclear membrane (INM) with increasing confinement. **(D)** Nuclear area to perimeter ratio (top) and nuclear invagination ratio (bottom) for increasing confinement. **(E)** Exemplary confocal images of cells expressing Lap2B-eGFP (inner nuclear membrane) for increasing confinement. **(F)** Curvature analysis of nuclear shape for 20 consecutive frames (t_lag_=10 s) for unconfined (suspension, top) and 7 μm confined nuclei (bottom). **(G)** Histogram of nuclear curvature for unconfined and 7 μm confined nuclei. **(H)** Relative cortical myosin II intensity with respect to cell and nuclear size increase for cells dissociated from embryos at high-oblong (3.3 hpf) and sphere stage (4 hpf) and cultured under similar confinement heights as indicated. **(I)** Exemplary confocal images of progenitor cells expressing Myl12.1-eGFP under 13 μm confinement dissociated from 4 hpf (top) and 3.3 hpf (bottom) embryos. **(J)** Relative cortical myosin II enrichment for interphase and mitotic cells under 7 μm confinement cultured in suspension (control) in the presence of 1 μM Nocodazole. Exemplary confocal images of progenitor cells expressing Myl12.1-eGFP in interphase (yellow asterisk) or during mitosis (cyan asterisk) under 7 μm confinement. **(K)** Exemplary confocal images of progenitor cells expressing Myl12.1-eGFP during mitosis treated with 1 μM Nocodazole and 50 μM LPA. Data points and error bars indicate mean and standard error of the mean (sem). ***p<0.001, **p<0.01, not significant (n.s.). All scale bars 10 μm despite 5 μm in (C).

To further probe the dependence of cortical myosin II accumulation on nucleus size, we dissociated primary embryonic stem cells from early and late blastula stages as cells reduce their size in consecutive rounds of early cleavage divisions (Fig. S3G). Deforming cells of different sizes under similar confinement heights revealed that myosin II accumulation is correlated with relative changes in nucleus deformation but not cell deformation (Fig. 2H). To test a functional role of the nucleus in regulating cortical contractility levels during cellular shape deformation, we analyzed cortical myosin II accumulation in mitotic cells that present a disassembled nuclear envelope. To arrest cells in mitosis and further increase the percentage of mitotic cells, we used Nocodazole. Confinement of mitotic cells (either spontaneous or Nocodazole-induced) did not trigger a cortical myosin II accumulation at 7 μm confinement height compared to interphase blastula cells (Fig. 2J), although they accumulated myosin II in response to LPA (Fig. 2K), a potent activator of Rho/Rock signaling, that has previously been shown to induce rapid cortical myosin II enrichment in zebrafish embryonic progenitor stem cells (22). Altogether, these data show that myosin II enrichment is correlated to nuclear shape deformation and stable INM membrane unfolding. This suggests that the nucleus functions as a continuous non-dissipative sensor element of cell deformation involved in the mechanosensitive regulation of cortical contractility levels and cellular morphodynamics.

To directly test biophysical nuclear characteristics, we developed an assay to probe intracellular nucleus mechanics by optical tweezer measurements. For this purpose, latex beads of 1 μm size were injected into 1-cell stage embryos that dispersed across embryonic cells during early cleavage cycles and acted as intracellular force probes to measure rheological properties of the nucleus (Fig. S4A). Trapezoidal loads were measured for cells in suspension and under 10 μm confinement (Fig. S4B-E). The recorded force followed the fast-initial indentation to reach a peak force before it relaxed to a non-zero constant force-plateau. The relaxation time remained unchanged between suspension (τ=3.14s +/-0.5s) and confined cells (τ =3.13s +/-0.6s) (Fig. S4D-H), suggesting passive but rapid (second scale) relaxation of a viscous component. The force-plateau on long timescale corresponds to an elastic component of the nucleus (Fig. S4I), in line with previous measurements that identified an elastic behavior of the nucleus (31) that can act as a cellular strain-gauge.

We next aimed at identifying nucleus deformation-dependent signaling pathways that link the spatio-temporal correlation of nuclear shape changes with fast myosin II activation and changes in morphodynamic cell behavior. Our previous observations suggested that nucleus deformation and associated mechanosensitive processes at the INM interface are involved in the regulation of myosin II activity and cortical contractility. Among a set of molecules tested under confinement conditions, we identified cytosolic phospholipase A2 (cPLA2) as a key molecular target mediating the activation of cortical myosin II enrichment under cell compression (Fig. 3A,B). Inhibition of cPLA2 by pharmacological or genetic interference robustly blocked cortical myosin II re-localization under varying confinement heights (Fig. S5A). Overexpression of cPLA2 mRNA rescued the morphant phenotype and led to a comparable myosin II accumulation as in control cells (Fig. 3A,B). To exclude that other mechanisms such as structural changes in the actin network prevent cortical myosin II re-localization under cPLA2 inhibition, we added LPA as an exogenous myosin II activator to cPLA2 inhibited cells. Under this condition, myosin II was strongly accumulated at the cell cortex (Fig. S5B,C) and induced cell polarization and amoeboid motility (Fig. 3C), suggesting that myosin II can be activated by alternative pathways when cPLA2 signaling is inhibited and is competent to bind to the cell cortex.

**Fig. 3.**
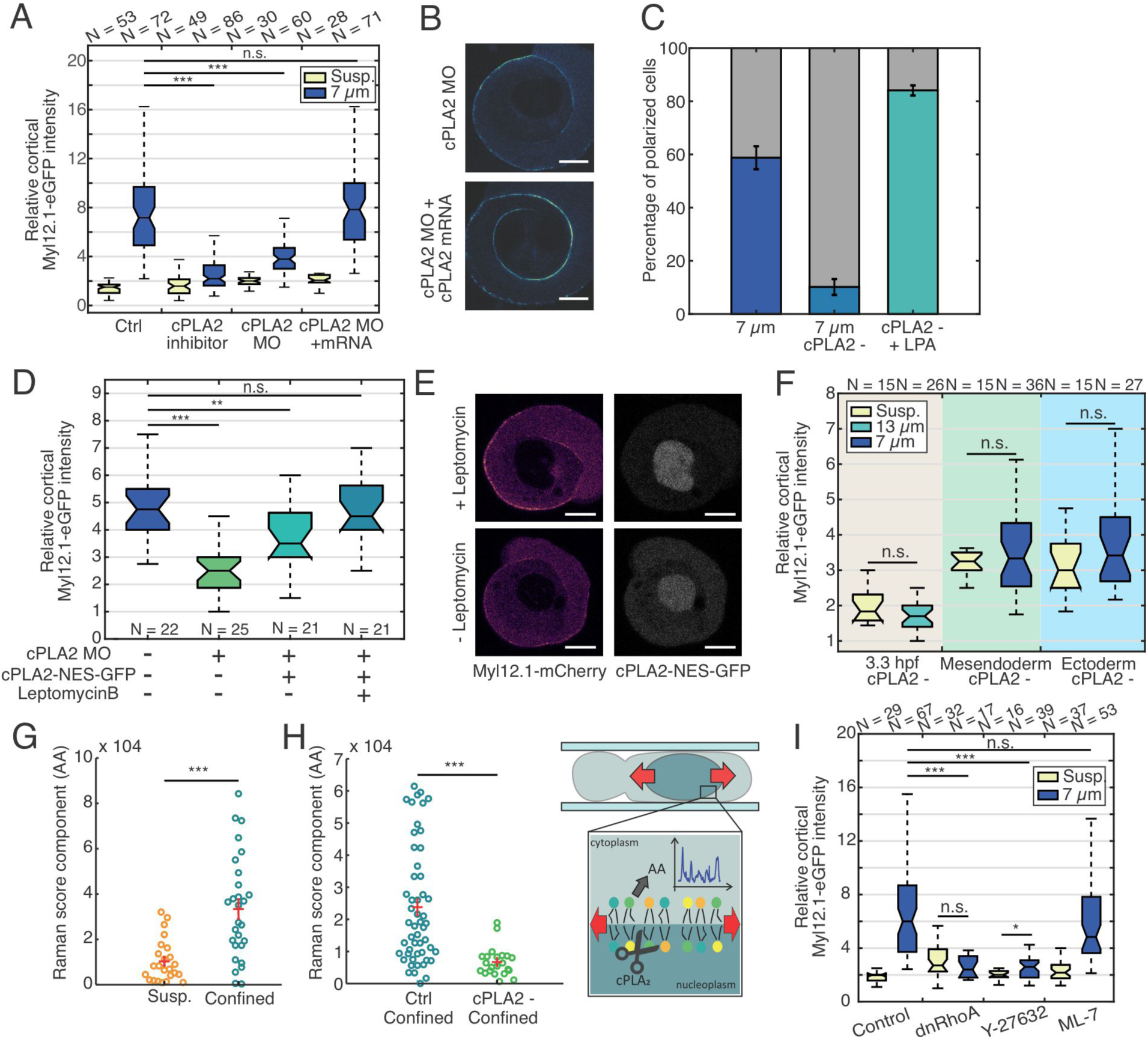
Nucleus deformation activates a mechanosensitive lipase signaling pathway regulating myosin II activity. **(A)** Relative cortical myosin II intensity for progenitor cells cultured in suspension versus 7 μm confinement conditions for control cells (DMEM), with cPLA_2_ inhibitor, or injected with cPLA_2_ MO and cPLA2 morpholino+cPLA_2_ mRNA. **(B)** Exemplary confocal images of progenitor cells expressing Myl12.1-eGFP under 7 μm confinement dissociated from embryos injected with cPLA_2_ MO (top) and cPLA_2_ morpholino+cPLA_2_ mRNA (cPLA_2_ rescue, bottom). **(C)** Percentage of stable-bleb polarized cells for control cells under 7 μm confinement, with cPLA_2_ inhibitor under 7 μm confinement or unconfined and stimulated with 50 μM LPA. For all conditions: n>200. **(D)** Relative cortical myosin II fluorescence intensity for cells dissociated from controls (un-injected) embryos or embryos injected with cPLA_2_ MO, cPLA_2_ MO+cPLA_2_-NES-GFP mRNA with or without Leptomycin B treatment. **(E)** Exemplary confocal fluorescence images of cell expressing Myl12.1-mCherry (left) and cPLA_2_-NES-GFP (right) under 7 μm confinement with (top) or without (bottom) the addition of Leptomycin B. **(F)** Relative cortical myosin II fluorescence intensity upon cPLA_2_ inhibition for cells dissociated at 3.3 hpf, induced mesendoderm or ectoderm cells in suspension and upon confinement at indicated height. **(G)** Scores of Raman component associated to AA in suspension (unconfined) and confined cells (10 μm). Red lines indicate mean and standard error of the mean (sem). **(H)** Scores of Raman component associated to AA in confined cells (10 μm) cultured in control condition or treated with cPLA_2_ inhibitor. Red lines indicate mean and sem. **(I)** Relative cortical myosin II intensity for control cells and different chemical (Y-27637, M-L7) or genetic interference (dnRhoA) with myosin II regulators. ***p<0.001, **p<0.01, *p<0.05, not significant (n.s.). All scale bars 10 μm.

Recent work identified that the activation of pro-inflammatory signaling during leucocyte recruitment to wounding sites is regulated by tension-sensitive binding of cPLA2 to the INM (32). We thus tested a role of cPLA2 in the nucleus by generating a modified cPLA2 construct containing a nuclear export sequence (NES). Using Leptomycin B as a blocker of nuclear export, an accumulation of cPLA2-NES-GFP within the nucleus was observed, showing a concomitant increase of cortical myosin II levels in confined cells (Fig. 3D,E). These data support, that cPLA2 localization in the nucleus is required for myosin II enrichment at the cortex.

We further validated that cortical myosin II enrichment in cells of different sizes (early versus late blastula cells) and different embryonic cell lineages (mesendoderm/ectoderm) depends on the activation of cPLA2 signaling. Pharmacological inhibition of cPLA2 activity blocked cortical myosin re-localization under cell deformation in confinement (Fig. 3F), supporting a consistent role of cPLA2 activation under physical cell deformation across early to late developmental stages. These data support that activation of cPLA2 signaling in the nucleus mediates adaptive cytoskeletal and morphodynamic behavior under cell deformation.

Arachidonic acid (AA) is the primary cleavage product generated by cPLA2 activity (33). To directly validate whether nucleus deformation in confinement triggers cPLA2 activity, we measured the release of AA by Raman spectroscopy. The analysis of Raman spectra confirmed the specific production of AA in confined cells (Fig. 3G and Fig. S5D), with the increase in AA production in confined versus control cells being specifically blocked in the presence of cPLA2 inhibitor (Fig. 3H). We further observed that AA was exclusively detected in the cytoplasm of confined cells, arguing that AA is directly released from nuclear membranes into the cytoplasm. These data support that cell confinement leads to enhanced cPLA2 activity and production of arachidonic acid associated with INM unfolding and stretching of the nucleus surface.

AA has been implicated to regulate myosin II activity both directly (34) and indirectly via protein phosphorylation (35). We tested the involvement of Rho/ROCK and Calcium-MLCK signaling that act as key regulatory pathways of myosin II activity (4). MLCK inhibition showed no significant effect on myosin II enrichment in confined cells, while a pronounced reduction of cortical myosin recruitment was observed under inhibition of Rho activity (Fig. 3I). Using a RhoAFret sensor further indicated an increased RhoA activity in confined cells versus control cells in suspension (Fig. S5E,F). These data support that AA production by cPLA2 activity engages upon nuclear envelope unfolding, regulating phosphorylation-dependent myosin II activity at the cell cortex.

Interference with intracellular calcium levels by addition of BAPTA-AM further induced a strong decrease of myosin II enrichment in confined cells, without altering cortical myosin II levels in unconfined control cells (Fig. S5G). LPA stimulation of BAPTA-AM treated cells confirmed that myosin II can be activated by the Rho-ROCK signaling pathway in the absence of intracellular calcium and remains competent to bind the cell cortex (Fig. S5B). Similarly, chelating extracellular calcium or using cPLA2 inhibition in combination with BAPTA-AM reduced cortical myosin II re-localization, while depletion of internal calcium stores via Thapsigargin led to a slight increase in myosin II enrichment in confinement (Fig. S5G). The addition of ionomycin showed that high intracellular calcium levels, in the absence of cellular shape deformation, were not sufficient to evoke AA production (Fig. S5H) and cortical myosin II enrichment (Fig. S5B). This suggests that intracellular calcium has a permissive function for increasing cortical contractility under cell confinement, in line with the observation that cPLA2 contains a calcium-dependent C2 binding domain that modulates protein activity.

To study if INM unfolding during nucleus deformation under cell confinement is sufficient to trigger cPLA2 activation, we performed a complementary experiment to induce cellular shape deformation by hypotonic swelling of cells. Quantification of nuclear shape parameters (size, volume, surface) revealed that hypotonic swelling induced comparable nuclear area expansion and INM unfolding as nucleus deformation under a confinement height of 7 μm (Fig. S6A,B). Interestingly, myosin II cortical enrichment in hypotonic conditions (Fig. 4A, Movie 7) and associated changes in bleb size (Fig. S6C) and cell polarization rate (Fig. 4C) were significantly lower compared to cells deformed at 7 μm confinement height. These observations suggest that nuclear envelope unfolding is not sufficient to trigger high levels of cortical myosin II enrichment under isotropic cell stretching in hypotonic conditions versus anisotropic cell deformation in confinement.

**Fig. 4.**
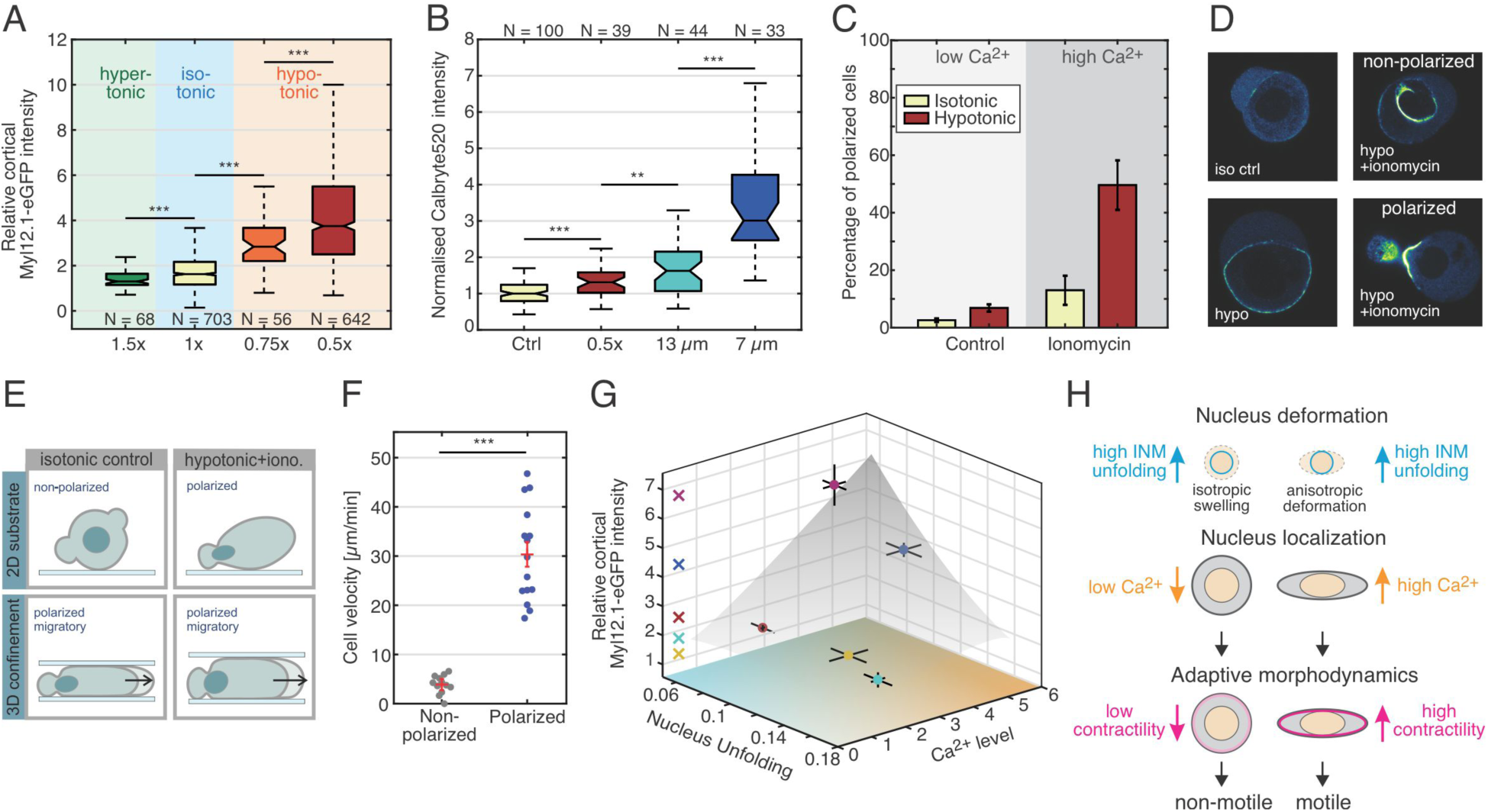
Nucleus unfolding and intracellular positioning enable adaptive cellular response to different types of physical cell deformation. **(A)** Relative cortical myosin II enrichment for progenitor cells cultured under different osmolarity conditions. **(B)** Normalized Ca^2+^ (Calbryte520) intensity for control (Ctrl) and hypotonic (0.5x) conditions and mechanical confinement (13 μm, 7 μm). **(C)** Percentage of stable-bleb polarized cells in isotonic and hypotonic (0.5x) conditions for cells cultured in DMEM (Control) or supplemented with 1 μM ionomycin. **(D)** Exemplary confocal images of cells expressing Myl12.1-eGFP in isotonic (ctrl, top-left), hypotonic (bottom-left) and hypotonic conditions supplemented with ionomycin treatment (right): non-polarized cell (top) and stable-bleb polarized cell (bottom). **(E)** Sketch of cell polarization and motile cell behavior in 2D (top) versus 3D confined environments (bottom) and control conditions (isotonic media; left) versus hypotonic conditions in the presence of ionomycin (right). (**F**) Instantaneous cell speed of non-polarized cells versus polarized stable-bleb cells migrating in 16 μm confinement. Red lines represent mean and standard error of the mean (sem). (**G**) Normalized relative cortical myosin II fluorescence intensity as a function of nucleus unfolding and normalized Ca^2+^ (Calbryte) intensity for different physical cell deformations (dark blue: 7 μm; light blue: 13 μm confinement; yellow: 7 μm confinement+Bapta-AM; red: hypotonic condition; magenta: hypotonic condition+ionomycin). Data indicate mean+/-sem. The gray area sketches the relation between cortical myosin II and nuclear deformation versus intracellular calcium levels. (**H**) Sketch depicting how nucleus deformation and intracellular nucleus positioning correlate with INM unfolding and intracellular calcium levels, that differentially regulate myosin II cortical contractility and cellular morphodynamics. ***p<0.001. Scale bars 10 μm.

Comparing intracellular calcium levels between deformed cells in confinement and under hypotonic conditions showed a pronounced difference in intracellular calcium levels that increased in confined cells (Fig. 4B). This observation suggests that the level of intracellular calcium can modulate morphodynamic cellular responses to different mechanical shape deformations. Of note, ectopically increasing intracellular calcium levels under hypotonic conditions via the addition of ionomycin led to a pronounced and rapid increase in cortical myosin II enrichment (Fig. S6D,E, Movie 7) that triggered spontaneous cell polarization (Fig. 4C,D and Fig. S6F), and transformed non-motile cells into a highly motile stable-bleb amoeboid mode with fast migration speed under confinement ex vivo and in vivo (Fig. 4E,F and Fig. S6G,H,J, Movie 7,8). This increase in myosin II accumulation and motile cell transformation was significantly reduced under conditions interfering with cPLA2 activity (Fig. S6D,F). Together, these data reveal that intracellular calcium levels are differentially regulated under distinct shape deformations: uniaxial compression in confinement induces high intracellular calcium levels, while isotropic radial stretch in hypotonic stress conditions does only lead to a minor increase of intracellular calcium levels. Independently modulating nucleus deformation and calcium levels under different shape deformations confirmed that both parameters engage synergistically to modulate cortical contractility (Fig. 4G), thereby enabling a cell to distinguish between different types of shape deformation and to acquire a specific morphodynamic response.

Intracellular nucleus positioning appeared as promising candidate to differentially modulate calcium levels. ER-plasma membrane (ER-PM) proximity has been implicated as an important regulator of cellular calcium signaling (36). Visualization of membrane proximal ER structures showed that the ER was highly dynamic under conditions of low confinement, but was increasingly immobilized between the nucleus-PM interface for larger cell deformations in confinement (Movie 9). In addition, the expanding nucleus contact area close to the plasma membrane closely correlated with an intracellular calcium increase (Fig. S6I). These observations suggest that mechanical compression induces a tight connectivity between ER-PM structures and that an increased ER-PM contact interface and/or mechanical stress within the ER directly triggers an intracellular calcium increase.

Together, our data suggest that the nucleus establishes a core element to measure cellular shape deformation via two key physical parameters: 1) nuclear shape deformation leading to INM unfolding and 2) intracellular spatial positioning of the nucleus. In this model, INM unfolding under nuclear shape change allows for the deformation-dependent activation of cPLA2 signaling, whereby cPLA2 activity is modulated by intracellular calcium levels set by nucleus-PM proximity (Fig. 4H, Fig. S7A). Our data reveal that the parameter space of these two variables (INM unfolding and calcium levels) provides a unique identifier for a cell to decode distinct shape deformations as exemplified on anisotropic cell deformation in confinement versus isotropic hypotonic cell stretching, allowing cells to acquire a unique adaptive response depending on the type of physical shape deformation (Fig. S7B).

Biochemical, physical and mechanical cues in the surrounding of a cell create manifold information for cells which is continuously sensed, integrated and transduced to allow for complex cellular functioning. Here we show that the cell nucleus functions as a cellular mechano-gauge for precisely decoding cellular shape changes, allowing cells to adaptively and rapidly tune cytoskeletal network properties and morphodynamic behavior within their 3D tissue microenvironment during development. This mechanism is one of the first to lay a foundation for functional principles underlying cellular proprioception that, comparable to the sensing of spatio-temporal changes in body posture and movement (37), enable a precise interpretation of shape changes on the single cell level. The nucleus, being the largest organelle within the cell, represents a prominent structure to transmit and modulate mechanosensitive processes, and has been shown to influence cell differentiation (38-40), migration (41-43) and pathfinding in constrained environments (44).

Our findings support that nucleus deformation and its intracellular positioning establish a cellular sensing module that equips cells to rapidly and reversibly adapt their dynamic response to shape fluctuations. Such a mechanism is relevant for various biological processes as migration plasticity of cancer and immune cells in constrained 3D tissue niches (45, 46), mechano-chemical feedback processes during morphogenesis (47) and homeostatic tissue functions (14), which require accurate mechanisms to detect variations in cellular size and shape and multicellular packing density in crowded 3D tissues.

## Acknowledgments

The authors would like to acknowledge the Super Resolution Light Microcopy and Nanoscopy (SLN) Facility of ICFO for their support with imaging experiments and Johann Osmond (Nanofabrication laboratory, ICFO) for the design and production of molds for generating confinement coverslip, and further support from the CRG Core Facilities for Genomics and Advanced Light Microscopy. We thank the following labs that kindly provided plasmids: pCS2+ lefty/casanova (courtesy Carl-Philipp Heisenberg); pCS2+ lyn-TdTomato (courtesy Berta Alsina); pTriEx-RhoA FLARE.sc Biosensor WT was a gift from Klaus Hahn (Addgene plasmid #12150; RRID:Addgene_12150). We would like to thank Carl-Philipp Heisenberg, Matthieu Piel and Alexis J. Lomakin for discussions on this work and Ben Lehner, Vivek Malhotra, Sebastian P. Maurer and the Ruprecht and Wieser lab members for critical reading of the manuscript.

## Authors contribution

V.R. and S.W. designed research; V.V. performed key experiments and data analysis; F.P. contributed to hypotonic and interference experiments and RNA preparations; F.C-C. and V.V. performed optical tweezers experiments and F.C. analyzed the data; M.M-S. and V.V. performed Raman experiments and M.M-S. analyzed the data; M.C-R. analyzed the Lap2B-GFP data; H-M.M. and Q.T.-R. performed in vivo experiments and H.-M.H. performed injections and helped with mesendoderm-ectoderm experiments; S.J-D. cloned plasmids, synthetized mRNA and performed mRNA and bead injections; S.P-L. performed calcium imaging related to the role of Piezo channels; M.A.V. supervised S.P-L. and contributed with discussions and support to calcium imaging experiments; M.K. supervised F.C.-C. and designed tweezer experiments. S.W. and V.R. supervised the project, contributed to data analysis and wrote the manuscript.

## Data and materials availability

All data is available in the main text or the supplementary materials.

