## Supplementary_Materials for "The nucleus measures shape deformation for cellular proprioception and regulates adaptive morphodynamics"

**This PDF file includes:**

Materials and Methods
Figs. S1 to S7
Captions for Movies S1 to S9

### Materials and Methods

**Zebrafish Maintenance** Zebrafish (Danio rerio) were maintained as previously described (*1*). Embryos were kept in E3 medium at 25°C–31°C prior to experiments and staged based on morphological criteria (*2*) and hours post fertilization (hpf). Wild-type embryos were obtained from the AB strain background. All protocols used have been approved by the Institutional Animal Care and Use Ethic Committee (PRBB–IACUEC), and implemented according to national and European regulations. All experiments were carried out in accordance with the principles of the 3Rs.

**Transgenic fish lines** The following transgenic lines were used: Tg(actb2:Lifeact-GFP) (*3*), Tg(actb2:Myl12.1-eGFP) (*4*), Tg(actb2:Myl12.1-mcherry) (*4*), Tg(actb2:Lyn-TdTomato) (*5*). All progenitor cells expressing Myl12.1-eGFP (Myosin II) and Lifeact-GFP (Actin) were obtained from Tg(actb2:Myl12.1-eGFP) and Tg(actb2:Lifeact-GFP).

**Cell Culture** To culture progenitor cells, embryos were manually dechorionated in E3 buffer at sphere stage (4 hpf) or different stages if indicated. Five to twenty embryos were transferred to DMEM-F12 (with L-Glut and 15mM HEPES, without sodium bicarbonate and phenol red) culture medium (Sigma) and mechanically dissociated by manual tapping followed by centrifugation at 200g for 3 min.

**Sample preparation and surface coatings** The following products for surface coatings at the indicated concentration have been used: 0.5mg/ml PLL(20)-g[3,5]-PEG(2) (Susos) and 0.2 mg/ml fibronectin (bovine plasma from Sigma-Aldrich). Prior to PLL-PEG coating, both coverslips and dishes were plasma cleaned. Uncoated or PLL-coated glass dishes #1.5 were purchased from MatTek (MatTek Corporation).

**Cell Confiner** Cells were confined using a dynamic confiner (4DCell) similar to previously established planar microconfinement methods (*6*). In order to confine cells at different heights, multiple Si molds were produced by photolithography in a clean room (Nanofabrication laboratory, ICFO) by depositing a SU-8 resin on a silicon wafer. In brief, a photomask with the desired geometry was created. Confinement coverslips were prepared with polydimethylsiloxane (PDMS) with the following heights: 18, 16, 13, 10, 7 and 5 μm. Coverslips were always plasma cleaned, coated with PLL-PEG and equilibrated in DMEM prior to each experiment. A pressure pump (AF1 – Microfluidic pressure pump, Elveflow) together with the ESI software was used to change the pressure for tuning the confinement heights. For Raman measurements and optical tweezers two coverslips separated with microbeads or with a PDMS membrane have been used (h=10 μm).

**Reagents and Inhibitor Treatments** Pharmacological inhibitors were used at the following concentrations: 1 μM $cPLA_2$ inhibitor (Pyrrophenone, Merck-Millipore), 10 μM Bapta-AM (cayman), 10 μM Blebbistatin (+) (Tocris Bioscience), 10 μM Y-27632 (Tocris Bioscience), 10 μM ML-7 hydrochloride (MLCK inhibitor, Tocris Bioscience), 1 μM Nocodazole (Sigma), 50 nM LeptomycinB (Sigma-Aldrich), 1 μM Ionomycin (Sigma-Aldrich), 1uM Thapsigargin (Thermofisher). 10μM GsMTx4 (Tocris), 100 μM Z-VAD(OMe)-FMK (caspase inhibitor, Abcam), 1-oleoyl lysophosphatidic acid (LPA, Tocris Bioscience) was used at the indicated concentrations. Measurements were done directly after exposure to MLCK inhibitor, Ionomycin, LPA; all the other inhibitors have been pre-incubated for 30 min and 60 min for Y-27632 prior to experiments.

**Fluorescence staining** Calbryte520 (AAT BIOQUEST) was used to study calcium dynamics. The staining kit-Red Fluorescence-Cytopainter (ER Tracker, Abcam) or ER-Tracker™ Green

(BODIPY™ FL Glibenclamide) were used to visualize the endoplasmic reticulum respectively for confocal 3D colors imaging and for TIRF microscopy experiment. DNA-Hoechst (Thermofisher) was used to stain the cell nucleus. Cells were incubated with Calbryte520 for 20 min, with ER-tracker for 30 min and with DNA-Hoechst for 7-10 min as reported in the corresponding protocols and at the indicated concentrations. After incubation, cells were washed, centrifuged at 200 g for 3 min and re-suspended in DMEM media.

**Variable osmotic culture conditions** D-Mannitol (Sigma) was diluted in DMEM in order to obtain a culture medium with an osmolarity of approx. 450 mOsm/l (corresponding to a 1.5x media). Milli-Q water was added to DMEM for hypotonic conditions.

**Plasmid cloning** The following constructs were subcloned in a pCS2+ vector, linearized with BamHI-EcoRI restriction enzymes, using Gibson cloning system. LAP2b-eGFP was amplified from a pME 18S-FL3 vector (clone #2643665, Dharmacon); cPLA2a (pla2g4aa) plasmid was amplified from a pCR4-TOPO vector (clone #9037889, Dharmacon). All cDNAs were amplified using Phusion HF DNA Polymerase (Thermofisher F530S), see primers below. NES-cPLA2a-eGFP was amplified from the pCS2+-cPLA2a-eGFP vector using primers encoding for an N-terminal NES sequence (LPPLERLTL).The following constructs were provided from different labs: pCS2-DNRhoA N19 (*7*); pCS2+_cyclops (*8*), pCS2+ lefty and Casanova (courtesy Carl-Philipp Heisenberg); pCS2+ Lyn-TdTomato (courtesy Berta Alsina); pTriEx-RhoA FLARE.sc Biosensor WT was a gift from Klaus Hahn (Addgene plasmid #12150; RRID:Addgene_12150). Oligonucleotides used for cloning:

| | |
|---|---|
| pCS2-LAP2b-eGFP Fw 1 | 5'-GCTACTTGTTCTTTTTGCAGGATCCATGTCGGAATTTCTGGAAGACCC-3' |
| pCS2-LAP2b-eGFP Rv 1 | 5'-CTATTGAGGGCTCAGAGAAATCCTTGT-3' |
| pCS2-LAP2b-eGFP Fw 2 | 5'-TTTCTCTGAGCCCTCAATAGTGAAGGAG-3' |
| pCS2-LAP2b-eGFP Rv 2 | 5'-CCTCGCCCTTGCTCACCATGAATTCTTTGCTGGTACTGTCATCTGTGCC-3' |
| pCS2-cPLA2a Fw | 5'-GCTACTTGTTCTTTTTGCAGGATCCGCCACCATGTCCAACATTATAgtaagtgacgcttG-3' |
| pCS2-cPLA2a Rv | 5'-GCTCGAGAGGCCTTGAATTCTCACACTTTTGTGTAGCTTTTTGCA-3' |
| pCS2_NES-cPLA2a-eGFP Fw | 5'-GCTACTTGTTCTTTTTGCAGGATCCGCCACCATGCTGCCCCCCCTGGAGCGCCTGACCCTGTCCAACATTATAGTTG -3' |
| pCS2_NES-cPLA2a-eGFP Rv | 5'- GCTCGAGAGGCCTTGAATTCCTAGAgCTTGTACAGCTCGTCC-3' |

**Zebrafish mRNA injections** mRNA was synthesized using the mMessage mMachine Kit SP6 Kit (Ambion AM1340M). All the mRNA injections were done in 1-cell stage embryos.

To visualize the inner nuclear membrane 80 pg of Lap2B-eGFP were injected in wild type AB or Tg(actb2:Lyn-TdTomato). To interfere with myosin II regulators 100 pg of dn-RhoA mRNA (*7*) have been injected in Tg(actb2:Myl12.1-eGFP) together with 100 pg LynTomato mRNA (to visualize plasma membrane). For the RhoA-Fret imaging 400p g of RhoA-Biosensor (*9*) were injected in wild type embryos. To induce mesendoderm, endoderm, mesoderm or ectoderm cells, 1-cell stage wild type or Tg(actb2:Myl12.1-eGFP) embryos were injected respectively with: 100 pg cyclops mRNA, 50 pg Casanova mRNA, 100 pg cyclops mRNA and 2 ng Casanova morpholino (GCATCCGGTCGAGATACATGCTGTT), 100 pg Lefty mRNA, all supplemented with 100 pg LynTomato mRNA.

**Morpholino interference and rescue experiments** For inhibiting cPLA2 activity, 2.7 ng of cPLA2 morpholino (AAGCGTCACTTACTATAATGTTGGA) were injected in 1-cell stage Tg(actb2:Myl12.1-eGFP) or Tg(actb2:Myl12.1-mcherry) embryos. To rescue the activity, cPLA2 morpholino was co-injected with 200 pg of cPLA2 mRNA and 100 pg of LynTomato mRNA or 200pg of NES-cPLA2-GFP mRNA. The same concentrations of mRNA were used for control experiments.

**Zebrafish Blastula injections** For injection of hypotonic media, Zebrafish embryos at sphere stage were dechorionated and placed in single embryo agarose wells (Adaptive Science Tools). Injections were applied into the extracellular space at the animal pole. Each embryo was injected with an average of 16 nl of injection mix containing hypotonic media (D3:MilliQ 1:1) with Dextran and Alexa Fluor™ 546 (10kMW, Anionic,ThermoFisher Scientific) to label the interstitial fluid and together with 10uM Ionomycin calcium salt (Sigma-Aldrich).

**Embryo mounting:** Embryos were mounted in 2% low melting point agarose prepared in Danieau's solution (58mM NaCl, 0.7mM KCl, 0.4mM MgSO4, 0.6mM Ca(NO3)2 and 5mM HEPES) on a Mattek dish and covered with Danieau's solution.

**Fluorescence imaging** Confocal fluorescence images were acquired using a commercial Leica TCS SP5 STED CW or Leica TCS SP8 STED 3X microscope equipped with a white light laser source (Leica Microsystems, Wetzlar, Germany). In both cases we used a 63x oil objective (NA=1.49, HCX PL APO CS 63.0x1.40 oil UV). For myosin II-eGFP, LifeACT-eGFP, Lap2B-eGFP and Calbryte520, $cPLA_2$-NES-GFP imaging, samples were excited with a 488nm light (Argon laser) using the SP5 microscope. For co-staining with the Lyn-Tomato membrane reporter or for the myosin II-mCherry experiments, a HeNe laser at 543 nm has been used for excitation and consecutives images have been acquired. In both color channels fluorescence was collected using single molecule detectors (SMD-HyD) in photon counting mode and transmission light was collected using a forward PMT. The Leica SP8 confocal microscope has been used for three color imaging. Myosin-GFP and ER-tracker Red have been excited respectively at 488 nm and 587 nm using the tuneable white light laser and the fluorescence light was collected using two backwords HyD detectors in photon counting mode while the DNA-Hoechst has been excited with a 405 nm semiconductor laser and fluorescence recorded using a PMT. The same microscope was used for the FRET imaging: the CFP was excited with the 405nm laser and the YFP at 512 nm using the white light laser and the emitted photons have been collected using the HyD detector in photon counting modes. Transmission light was collected using a forward PMT. A temperature controller set at T=28.5°C was used for all the experiments.

**In vivo confocal imaging** Embryos were imaged using a Leica SP5 confocal microscope equipped with a Leica 20X NA 0.7 immersion objective using a 488 nm laser and the emitted fluorescence was detected with a HyD detector. The temperature during imaging was kept constant at 28.5°C using a temperature chamber.

**Bright field imaging** Bright field movies were acquired using a Leica DMI-LED microscope equipped with IDS-CMOS cameras (UI-3880LE-M-GL) and a Leica 0.4x C-mount. Air objectives

10x (NA=0.25) or 20x (NA=0.40) were used to image cell dynamics. Acquisition was controlled using µManager (*10*).

**Optical trapping experiments** The optical tweezer platform (SensoCell, Impetux Optics, Spain) consists of a continuous wave laser (λ=1064 nm, 5 W nominal output power, Azur Light) steered with a pair of acousto-optic deflectors mounted around an inverted research microscope (Nikon Eclipse Ti2) equipped with a spinning disk confocal microscope (Andor DragonFly 502) on top of an active isolation table (Newport). The laser is directed onto a microscope objective (MO, 60x/NA=1.2, water immersion, Nikon) after being expanded with a telescope to fill the MO entrance pupil, through the epi-fluorescence port. A short-pass dichroic mirror reflects the IR trapping beam and transmits both the excitation and emission light for fluorescence microscopy, as well as bright-field. The AOD-driven positioning conversion factors [MHz/µm] were calibrated from bright-field images of a trapped bead in water, drawn over a ±30 x ±30 µm region in the field of view. Beads were tracked with CISMM's Video Spot Tracker (http://cismm.web.unc.edu/). Trapping forces were measured from detecting light-momentum changes of the forward scattered light through an NA = 1.4, oil immersion condenser lens using in a commercial force sensor (Lunam, Impetux Optics, Spain). This allowed us to measure forces beyond the linear trapping regime, thus covering the full spectrum until ~300 pN, and allowing to work with lower laser powers during the in-vivo force measurements, as compared to standard back focal plane interferometry (*11,12*). To perform intracellular trapping, we injected a 0.5 nL drop of diluted (1:5) 1 µm polystyrene beads (Sigma-Aldrich) into the one-cell embryo. Cells were seeded in home-made trapping microchambers consisting of a bottom-dish (Wilco Glass, #1.5) and a 1 x 1 inch cover glass spaced with a double scotch, 90 µm high tape. The bottom dish was coated with Concanavalin A (0.05 mg/ml, 30 min, Sigma-Aldrich) to partially adhere the cells and avoid movement due to cellular blebbing. For the optical trapping experiments in confinement conditions, we used two cover glasses (60 x 24 cm, # 1.5, Ted Pella) to enclose a few µl of cells and 10µm PS beads used as spacers. Only cells containing one bead were used for each measurement. The bead was then trapped by manually directing the focused laser beam at 300 mW power (at the sample plane; 2.5 W output power) and the force-displacement cycles were applied by addressing the AODs and PSD with custom made software (LabView, National Instruments). The bead was then brought against the nuclear membrane while force was recorded, by addressing the AODs and PSD with custom software based in LabView.

After fast indentation perpendicular to the nuclear membrane (~2-3 µm), the trap position was kept constant for 10 s. The same was applied with no bead and cell obstructing the beam path to account for initial momentum changes arising from dynamic trap location. Upon indentation, an elastic increase in force was followed by a force relaxation that was fitted by the following expression: $f(t) = A + (B - A)t^{-p}e^{-t/\tau}$; where A [pN] is the static force given by the residual stress applied onto the nuclear membrane; B [pN] is the peak force; p is the exponent for the initial, short time scale power law decay; and τ [s] is the characteristic time for the exponential decay at longer time scale. τ agreed to that obtained from a linear fit onto the data plot at semi-logarithmic scale, as $\tau = -1/m$, where m is the slope of the fitted line. Normalized force relaxation profiles (Suppl. Fig. S4 F,G) were obtained by scaling the force as $(f(t) - A)/(B - A)$. τ was obtained for N = 6 cells in suspension and N = 7 cells under mechanical confinement. Data was processed with custom Matlab scripts.

Nuclear deformation was imaged during the trapping routine using a Nipkow spinning-disk confocal imaging platform (Nikon Eclipse Ti2). To block the trapping laser light, an IR filter (IR-

F) was placed in the beam path of the microscope. Nuclear DNA was stained after incubation for 5-10 min in 5 μg/μl Hoechst dye and cells were excited with λ = 405 nm (DNA-Hoechst staining) and λ = 488 nm (myosin II-eGFP). The two laser lines were transmitted through a multi-band dichroic (D2, 405-488-561-637 nm, AHF) to simultaneously excite the two fluorophores in the sample. After emission, light is reflected by D2 and directed into a long-pass dichroic (500 nm), which enables parallel imaging of the two channels using two back-illuminated sCMOS cameras (Sona, Andor) after passing through emission filters 445/46 nm and 521/38 nm. Image acquisition was performed using Fusion Software and post-processed in Fiji. Tracking of nuclear morphodynamics was carried out by fitting a double sigmoid function over the segment along the indentation direction with a custom script in Matlab (Mathworks).

**Raman Spectroscopy (RS)** For RS experiments, progenitor cells dissociated from wild type embryos at sphere stage were used and measurements were carried out using re-suspending cells in DMEM media solution. For confinement conditions, a drop of cells was placed between two quartz coverslips (ESCO products, Oak Ridge, NJ). For non-confined cells, a separation of around 100 μm was left between the two coverslips. A total of minimum 25 spectra were obtained from different cells for each condition in the cytoplasm or nucleus. The Raman system (inVia Renishaw, Apply Innovation, Gloucestershire, U.K.) comprises a 532 nm laser (~10 mW) which is focused onto the sample plane using a 60X water immersion Nikon objective (backscattered configuration). The laser spot size is set to 0.8 μm allowing for localized Raman measurements. Raman spectrum were recorded on a deep depletion charge coupled device (CCD) detector (Renishaw RenCam). The recorded Raman spectrum is digitalized and displayed on a PC using Renishaw WiRE software. The spectra are background subtracted with a custom-written Matlab code (see methods in (*13*)). First, an exploration of the spectral data set was performed using Principal Component Analysis (PCA). Second Multivariate Curve Resolution (MCR) algorithm were performed to extract molecular components from the Raman spectral dataset (spectral profile and abundance in each measured sample). For PCA and MCR analysis, the PLS toolbox in Matlab was used.

**Calcium imaging (related to Suppl. Fig. 3D)** Progenitor cells were added to a Concanavalin A-coated (0.05 mg/mL by incubation at 31ºC for 1.5h) glass bottom dishes and loaded for 20 min at 28ºC with $5\,\mu M$ of FURA-2 plus an nonionic surfactant (0.02% pluronic F-127) dissolved in DMSO. The cells were then washed before initiating the experiment. The fluorescence signal was measured in a standard bath solution containing 140 mM NaCl, 2.5 mM KCl, 1.2 mM $CaCl_2$, 0.5 mM $MgCl_2$, 10 mM HEPES, and 5 mM glucose, pH 7.4, adjusted with NaOH ($\sim 320 - 340\, mOsm/L$). Video microscopic measurements of intracellular calcium concentrations were obtained using an Olympus IX70 inverted microscope (Hamburg, Germany) with a 40x oil-immersion objective (Olympus). A Polychrome IV monochromator (Till Photonics, Martinsried, Germany) supplied the excitation light (340 and 380 nm), which was directed toward the cells in the field of view by a 505DR dichromatic mirror (Omega Optical, Brattleboro, VT). Fluorescence images were collected by a digital charge-coupled device camera (Hamamatsu Photonics, Hamamatsu City, Japan), after their passage through a 535DF emission filter (Omega Optical), using the AquaCosmos software program (Hamamatsu Photonics). Cytosolic calcium concentration was presented as the ratio of emitted fluorescence after excitation at 340 and 380 nm relative to baseline.

**Data Analysis**

**Myosin/Actin accumulation and relative enrichment at the cortex** Myosin and actin relative cortical accumulation values in vitro are quantified from confocal images, acquired as described before, using a custom-written script in Matlab (2017b, Mathworks). The relative cortical accumulation is defined as: $(I_{cortex} - I_{bleb})/I_{cortex}$. Cortical intensities are quantified by manually selecting the cortical region where the fluorescence signals are homogenous. The statistics over many cells is used to calculate the median of the mean peak intensities of the cortical regions. The bleb intensity in a single cell is calculated by computing the mean intensity in a manually selected rectangular region in the bleb.

Relative myosin accumulation in vivo was quantified on raw data using Fiji (plot profile tool, Fiji) and for each ratio (peak to averaged cytosolic intensity fluorescence) the distance of the corresponding cell to the yolk margin was measured. Nuclear aspect ratio was calculated in Fiji (measurement tool, Fiji) and for each nucleus the normalized distance to the yolk margin was measured.

**Cell size, Bleb size and nuclear size estimation** Cell and nuclear diameter measurements are calculated with Fiji (Measure tool, Fiji) from Myosin II GFP confocal images; transmission images are used as control. Bleb sizes are quantified from bright fields movies by segmenting single cells and manually selecting the bleb and cell regions using Fiji (Measure tool, Fiji).

**Lap2B-eGFP nuclear analysis** Nuclear properties were analyzed using the LapB2-GFP images in Python using the Scikit-Image library (https://scikit-image.org/). We first determined the inner and outer outlines using the contour detection function and used a median filter for smoothing. From that we determined area, perimeter and the convex image. The ratio in between the inner area ($A_{in}$) and the associated convex area ($C_{in}$) is defined as the invagination ratio, computed as $= 1 - (A_{in}/C_{in})$ .

**Lap2B-eGFP shape analysis** Nuclear envelop fluctuations and bending analysis was done on Lap2B-eGFP cross-sectional confocal images. The nuclear envelop was manually tracked in Fiji and the discrete x-y positions were further post-processed using a custom made Matlab script. Between each pair of discrete x-y positions a cardinal spline function was interpolated (tension=0) passing through all x-y positions. The resulting spline vector was overlaid to the fluorescence image to manually control the match with the nuclear envelope circumference. The spline was further used to calculate the curvature along the line using 2D bending vectors. The histogram of bending vectors of 10 cells were compared between 7 μm confined cells and cells in suspension.

**Calcium imaging** Progenitor cells derived from WT embryos were stained with Calbryte520 a calcium activity reporter. The cell perimeter was segmented using either Myosin-mCherry, LynTomato or with a transmission image. The mean intensity in the Calbryte channel has been measured in Fiji.

**Percentage of polarized cells** Percentages of polarized cells were computed from bright field movies as number of polarized cells/total cells. Stable-bleb polarized cells can be easily distinguished from non-polarized cells due to their different morphology (pear shaped versus rounded blebbing).

**FRET** RhoA-FRET images have been analysed with the Fret analyser plugin in Fiji. Donor, acceptor and FRET images are used by the plugin to compute the FRET index and to obtain the FRET image.

**Statistical tests** Statistical significance test has been done with the two-sample t-test, suing the ttest2 function in Matlab. Datasets have been considered non-significant (n.s.) if $p>0.05$, and the following significance symbols have been used for the corresponding p-values: * $p<0.05$, ** $p<0.01$ and *** $p<0.001$.

### Supplementary Figures S1-S7

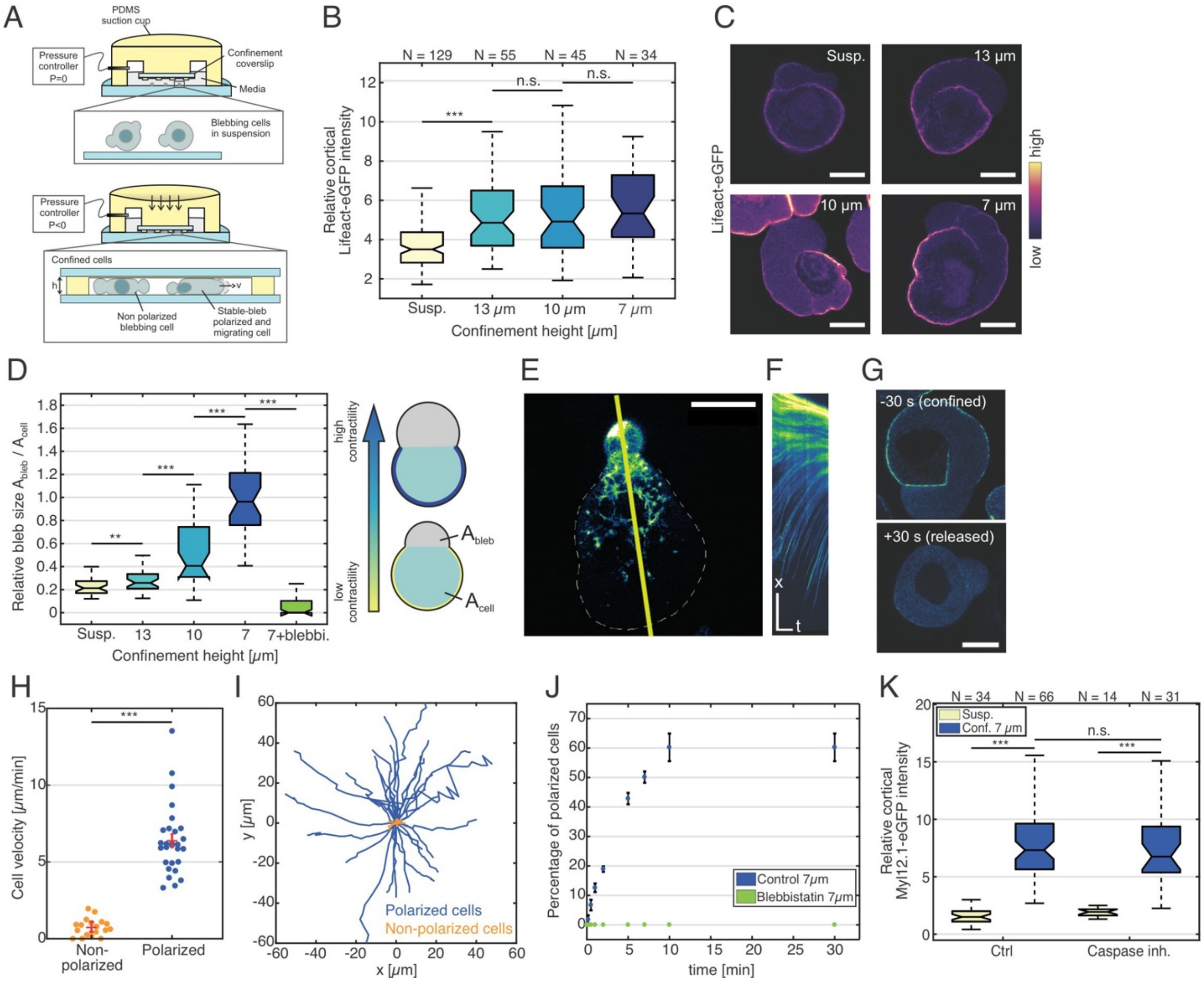


**Fig. S1.** (**A**) Sketch of plan-parallel cell microconfinement using coverslips with PDMS micro-pillars of desired heights, with plate distance regulated by a pressure controller. (**B**) Relative cortical LifeAct-GFP intensity for increasing confinement in un-polarized progenitor stem cells. (**C**) Exemplary confocal images of progenitor cells expressing Lifeact-GFP under different confinement heights. Scales bars 10 μm. (**D**) Normalized bleb to cell area for cells cultured in control conditions (Suspension, Susp.), different confinement heights and for 7 μm confinement supplemented with 10 μM Blebbistatin, for all conditions N=35. (**E**) Exemplary confocal image of the basal cortex of a stable-bleb cell expressing Myl12.1-eGFP under 7 μm confinement. Scale bar 20 μm. (**F**) Associated kymograph along yellow line in (E) showing retrograde cortical flow opposite to the direction of cell migration. Scale bars 10 μm (x) and 10 s (t). (**G**) Representative fluorescence image of a progenitor stem cell in 7 μm confinement before release of cell compression (-30 s) and after confinement release (+30 s). Scale bars 10 μm. (**H, I**) Mean cell velocity (H), and cell tracks (I) for polarized (blue) and non-polarized (orange) progenitor stem cells cultured in 7 μm confinement. Red lines represent mean and standard error of the mean. (**J**) Percentage of polarized migratory stable-bleb cells over time after applying 7 μm confinement (t=0) for progenitor stem cells cultured in control conditions (DMEM, blue) or with Blebbistatin (green); $t_{1/2}$~4min. (**K**) Relative cortical myosin II intensity for control cells or cells treated with caspase inhibitor in suspension and 7 µm confinement. ***p<0.001, **p<0.01, not significant (n.s.).

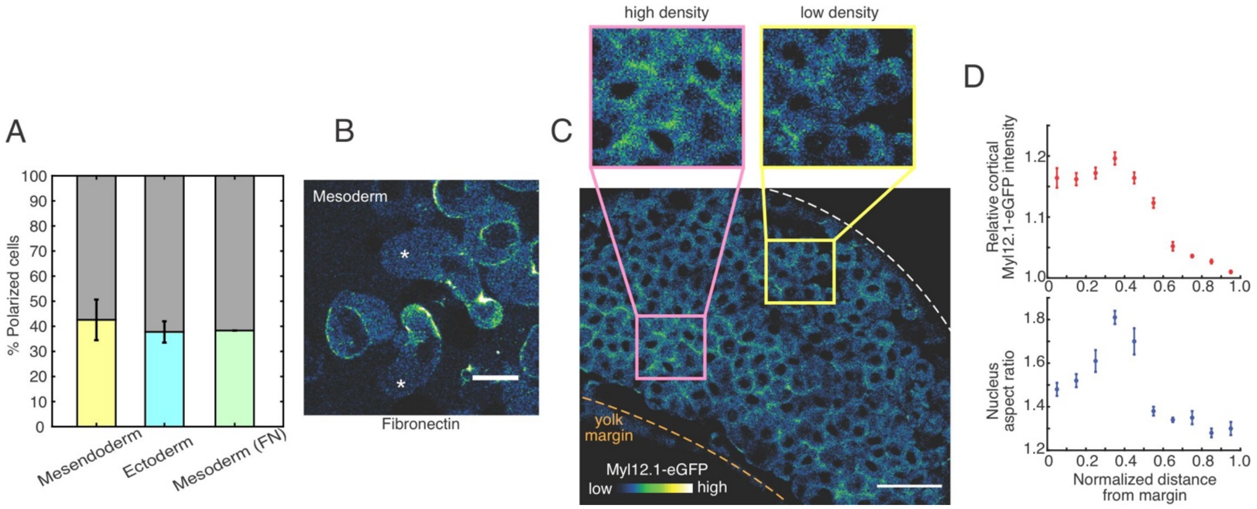


**Fig. S2.** (**A**) Percentage of cell polarization for induced mesendoderm and ectoderm cells on passivated surfaces and fibronectin-coated surfaces for 7 µm confinement. (**B**) Exemplary confocal images of induced mesoderm progenitor cells expressing Myl12.1-eGFP on a fibronectin-coated surface under 7 µm confinement. Asterisks indicates stable bleb polarized cells. Scale bar 20 µm. (**C**) Exemplary in vivo image of embryonic Myl12.1-eGFP distribution at the lateral margin at 4.5 hpf. The yolk interface (orange dashed line) and embryonic tissue surface (white dashed line) are highlighted. Scale bar 50 µm. **(D)** Relative cortical myosin II intensity and nuclear aspect ratio as a function of normalized distance from the yolk margin.

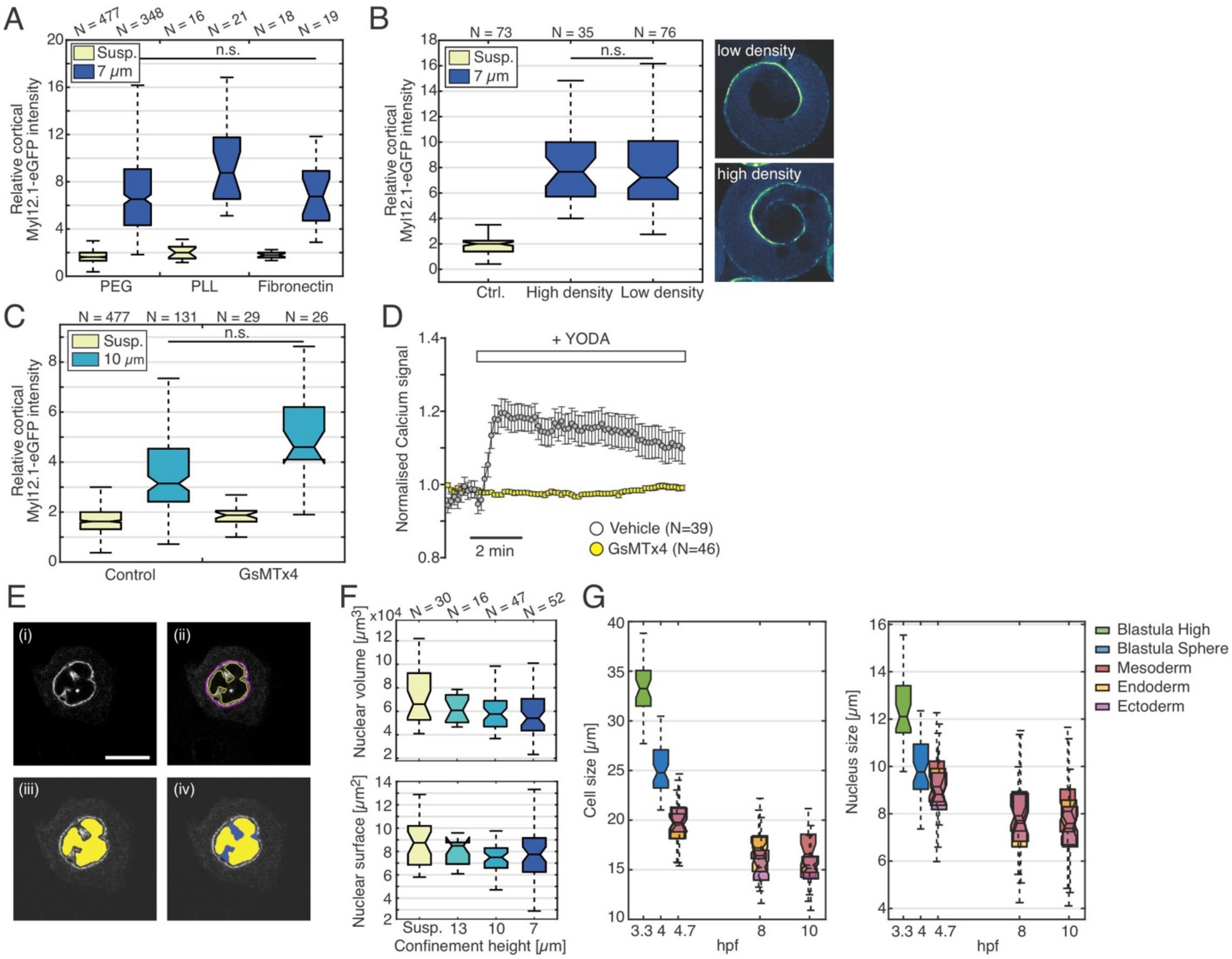


**Fig. S3.** (**A**) Relative cortical myosin II enrichment under 7 µm confinement for cells cultured on passivated (PLL-PEG) or adhesive surfaces (PLL, fibronectin). (**B**) Relative cortical myosin II intensity under 7 µm confinement for isolated cells (no cell-cell contact) or contacting cells under high cell density. Exemplary confocal images of progenitor cells expressing Myl12.1-eGFP under 7 µm confinement in low (top) or high-density (bottom) environment. (**C**) Relative cortical myosin II intensity for cells treated with the cationic mechanosensitive channels inhibitor GsMTx4 for cells under 10 µm confinement versus control cells in suspension. (**D**) Normalized calcium signal in isolated progenitor cells under stimulation with the Piezo1 activator YODA (gray) and in the presence of YODA supplemented with GsMTx4 (yellow). (**E**) Exemplary Lap2B-eGFP image analysis protocol: (i) raw image; (ii) detection of inner and outer contour (magenta and yellow line); (iii) inner surface area (yellow); (iv) detection of convex area of (iii), (blue). The invagination ratio (IR) is defined as 1 – the ratio in between the yellow and blue area. (**F**) Nuclear surface and volume for increasing confinement. (**G**) Cell size (left) and nuclear size (right) during embryo development from 3.3 hpf until 10 hpf for undifferentiated progenitor stem cells (3.3, 4hpf) and induced mesoderm, ectoderm and endoderm progenitor cells (4.7, 8, 10 hpf). All scale bars 10 µm. Not significant (n.s.).

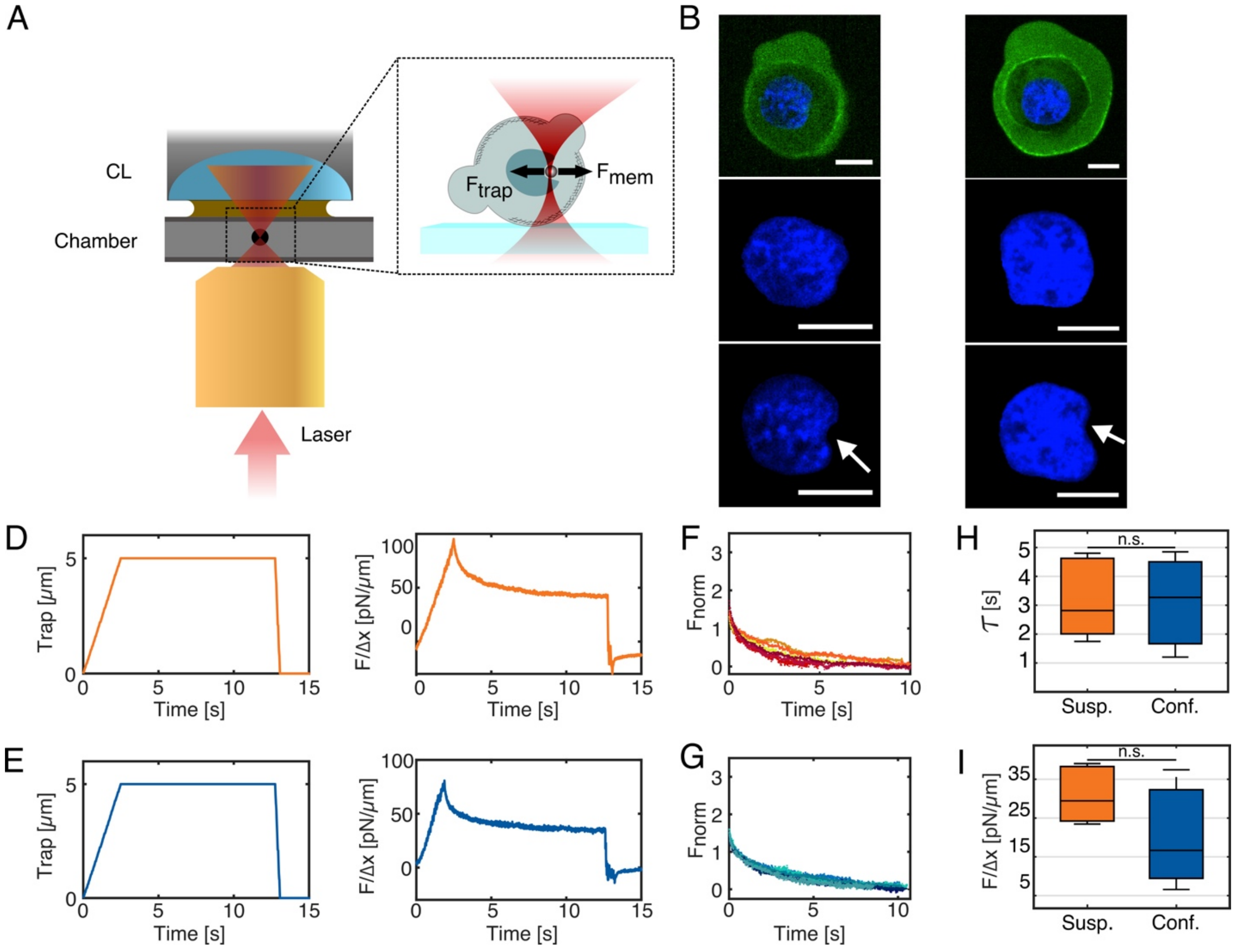


**Fig. S4.** Mechanical characterization of cell nuclei in live cells with optical micro-manipulation and spinning-disk confocal microscopy. (**A**) Schematic showing the trapping beam focused through a microscope objective and captured by a condenser lens (CL) to carry out light-momentum force measurements on an optical trap deforming the nuclear membrane. $F_{mem}$ and $F_{trap}$ are the nuclear membrane and trapping forces exerted onto the trapped 1 µm latex bead, respectively. **(B,C)** Exemplary fluorescence image of a cell in suspension (B) or confinement (C) expressing Myl12.1-eGFP and DNA-Hoechst (top) and snapshot prior to (middle) and during (bottom) indentation on the nucleus. The white arrow indicates the position of the trapped micro-bead. **(D,E)** Trap trajectory (left) and nuclear force profile (right, normalized to nuclear indentation) for exemplary force relaxation experiments in cells cultured in DMEM in suspension (D) or under 10 µm confinement (E). **(F,G)** Normalized and time-shifted nuclear force-relaxations tracks (as in panel D,E) for cells in suspension (F, N=6) and under 10 µm confinement (G, N=7). **(H)** Boxplot of characteristic time (τ) obtained from fitting an exponential decay function with plateau offset to the force-relaxation curves shown in (F,G). **(I)** Boxplot of static force component obtained from fitting an exponential decay to the force-relaxation curves. Not significant (n.s.).

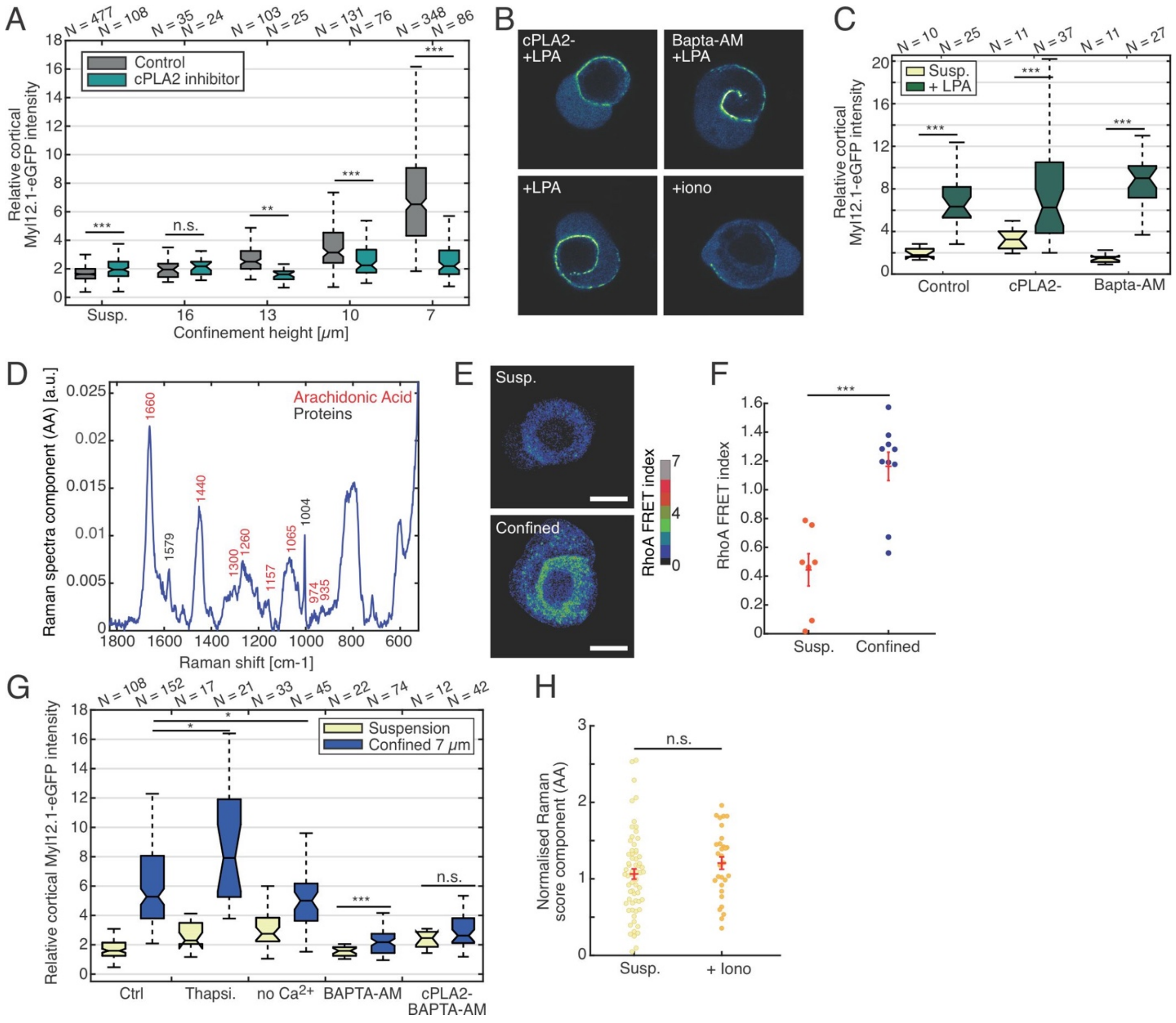

**Fig. S5.** (**A**) Relative cortical myosin II enrichment for cells cultured in suspension (DMEM) or supplemented with $cPLA_2$ inhibitor for decreasing confinement heights. (**B**) Exemplary confocal images of progenitor cells expressing Myl12.1-eGFP cultured in suspension with $cPLA_2$ inhibitor+LPA, Bapta-AM+LPA, LPA alone or ionomycin (iono). Scale bars 10 µm. (**C**) Relative cortical myosin II intensity for cells cultured in suspension (DMEM) and upon LPA addition in control conditions and for cells treated with $cPLA_2$ inhibitor or Bapta-AM. (**D**) Components of the Raman spectra associated with arachidonic acid (AA) used for the quantification of AA production (see Fig. 2). Raman peaks indicated in red are specific for AA. (**E**) Representative images showing the Fret index of control cells in suspension (Susp., unconfined) and 7 µm confinement. Scale bars 10 µm. (**F**) Scatter plot of FRET index for unconfined (suspension) and confined cells. (**G**) Relative cortical myosin II intensity for cells cultured in control (DMEM) or under different conditions, Thapsigargin (Thapsi.), Calcium free media (no $Ca^{2+}$), Bapta-AM and Bapta-AM+$cPLA_2$ inhibitor. (**H**) Scores of Raman component associated to AA in control (DMEM, suspension) and in the presence of 1 µM ionomycin (Iono). Red lines represent mean and standard error of the mean (sem). Red lines represent mean and (sem). ***p<0.001, **p<0.01, *p<0.05, not significant (n.s.).

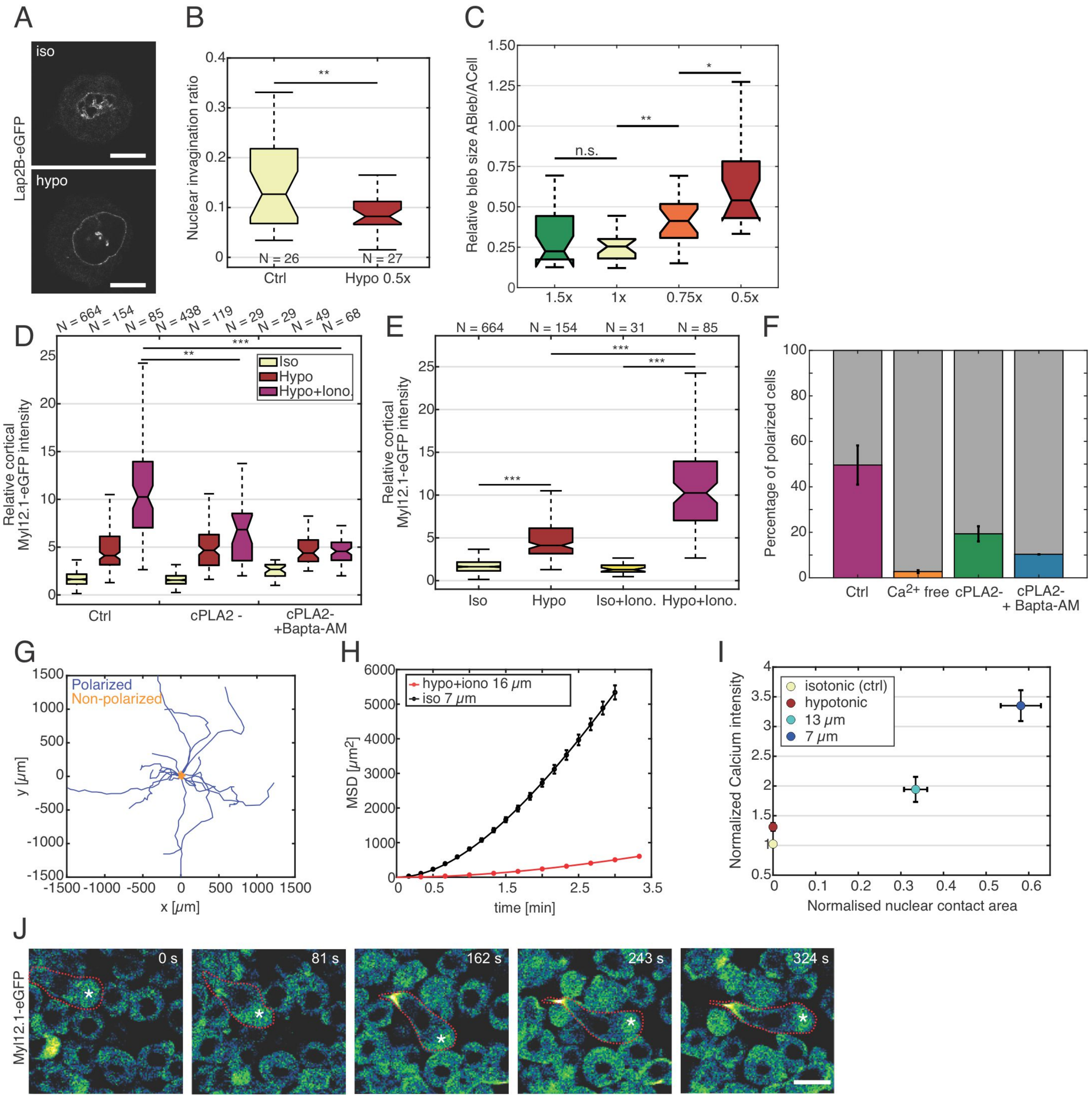


**Fig. S6.** (**A**) Exemplary confocal images of progenitor stem cells expressing Lap2B-eGFP cultured in isotonic (top) and hypotonic (bottom) conditions. Scale bars 10 μm. (**B**) Boxplot of nuclear invagination ratio for cells cultured in isotonic and hypotonic media. (**C**) Relative bleb to cell size under different osmolarity conditions, N=25 for each condition. (**D**) Relative cortical myosin II intensity for progenitor cells cultured in isotonic, hypotonic and hypotonic+ionomycin conditions supplemented with cPLA$_2$ inhibitor alone or in combination with Bapta-AM. (**E**) Relative cortical myosin II intensity for progenitor cells cultured in isotonic and hypotonic conditions and with ionomycin (Iono.). (**F**) Percentage of polarized stable-bleb cells under reference hypotonic conditions with 1 μM ionomycin (purple) and depletion of extracellular Calcium (orange), cPLA$_2$ inhibition (green) and cPLA$_2$+Bapta-AM (blue). (**G**) Cell tracks for polarized (blue, motile) and un-polarized (orange, non-motile) progenitor stem cells cultured in hypotonic media supplemented with 1 µM ionomycin under 16 µm confinement. (**H**) Mean square displacement (MSD) analysis of cell tracks related to (G). A persistent random walk model is fit to the data (Fürth formula) for 7 µm confined cells (black points) and hypo+ionomycin treated cells under 16 µm confinement (red points), with persistence time of $P_t$=1.9 min (2.8 min), respectively. (**I**) Mean calcium intensity with respect to the normalized nuclear contact area for isotonic condition (control, yellow), hypotonic shock (red), 13 µm confinement (light blue) and 7 µm confinement (dark blue). The nuclear contact area is normalized to the cross-sectional area for each cell. (**J**) Representative in vivo image of a motile stable-bleb cell (dashed red line) in a zebrafish embryo at blastula stage (4 hpf) injected with hypotonic media supplemented with ionomycin (10 µM). Asterisk denotes cell front. Data represent mean +/- sem. ***p<0.001, **p<0.01, *p<0.05, not significant (n.s.).

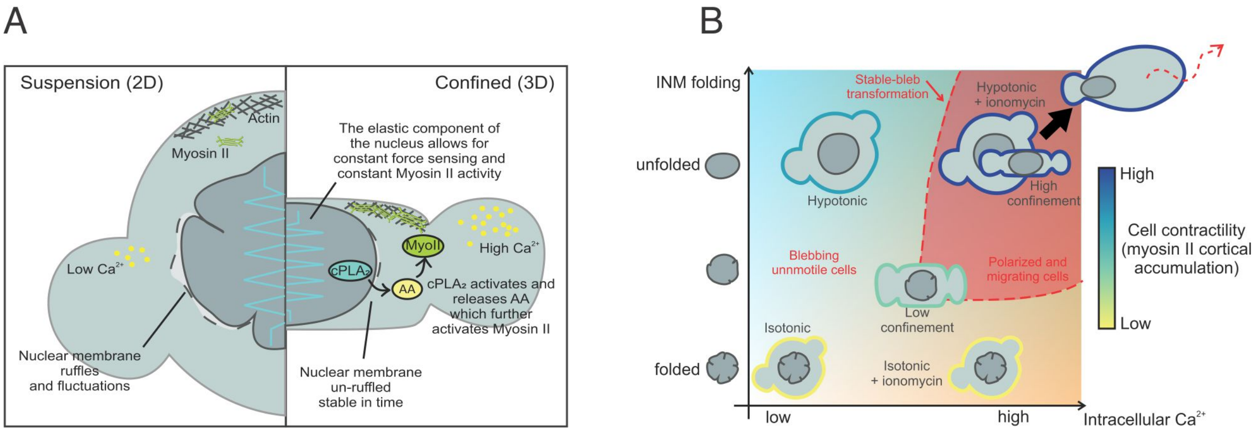


**Fig. S7.** (**A**) Illustration of the mechano-transduction pathway that translates nucleus deformation into myosin II re-localization and an increase in cortical contractility regulating morphodynamic migration plasticity. (**B**) Schematic representation of inner nuclear membrane unfolding and intracellular calcium levels depending on the type of physical cell deformation under anisotropic cell compression (top to bottom) and hypotonic swelling leading to isotropic cell stretching (left to right).

### Captions for Movies S1 to S9

**Movie S1.** Adaptive Myosin II dynamics upon cell confinement and role of $cPLA_2$ in cell mechanotransduction (related to Fig. 1 and Fig. 3). Time lapse confocal fluorescence movies of progenitor cells expressing Myl12.1-eGFP (myosin II) on non-adhesive substrates (PLL-PEG) cultured in DMEM media in suspension (left), under 7 μm confinement in control condition (middle) or supplemented with 1 μM $cPLA_2$ inhibitor. Upon confinement, myosin II accumulates at the cortex in control condition and the accumulation is blocked by the inhibition of $cPLA_2$.

**Movie S2.** Cortex remodelling during embryonic progenitor stable-bleb cell transformation (related to Fig. 1). Time lapse confocal fluorescence movie of Myl12.1-eGFP (myosin II) localization during stable bleb transformation of a progenitor cells under 7 μm mechanical confinement.

**Movie S3.** Myosin II retrograde flow in a stable-bleb polarized cell (related to Fig. 1 and Suppl. Fig. 1). Time lapse confocal fluorescence movie of the basal cortex of a stable-bleb polarized cell expressing Myl12.1-eGFP (myosin II) under 7 μm mechanical confinement. Myosin II shows a cortical density gradient and retrograde flow (opposite to migration direction).

**Movie S4.** Myosin II accumulation is reversible (related to Fig. 1). Confocal fluorescence time lapse movie of progenitor cells expressing Myl12.1-eGFP (myosin II) during confinement (10 μm) and subsequent release. Myosin II is rapidly re-localized from the cortex to the cytoplasm.

**Movie S5.** Mesenchymal-to-amoeboid transition: mesoderm cells transform to stable-bleb cells upon mechanical confinement (related to Fig. 1). Time lapse confocal fluorescence movie of mesendoderm-induced progenitor cells expressing Myl12.1-eGFP (myosin II) plated on fibronectin substrate showing mesenchymal migration (white asterisk indicates migrating cell, white arrow points at cell protrusions). Upon confinement (7 μm), cells rapidly accumulate myosin II at the cortex and transform to stable-bleb polarized cell (magenta asterisks point at cell front).

**Movie S6.** The inner nuclear membrane is unfolded under mechanical shape deformation in confinement and the unfolding is stable over time (related to Fig. 2). Time lapse confocal movies of progenitor cells stained with Lap2b-eGFP (i) in suspension, (ii) while confining cells at 7 μm and (iii) under 7 μm confinement.

**Movie S7.** Adaptive Myosin II dynamics under hypotonic conditions or ionomycin treatment (related to Fig. 4). Confocal fluorescence time lapse movie of progenitor cells expressing Myl12.1-eGFP (myosin II) on non-adhesive substrate (PLL-PEG) in isotonic media (control, DMEM), upon hypo-tonic shock (adding milliQ water, 0.5x shock) and upon the addition of 1 μM ionomycin. Hypotonic treatment is followed by a myosin II accumulation at the cortex; further ionomycin addition leads to a pronounced myosin II accumulation and triggers cell polarization.

**Movie S8.** Cell dynamics under hypotonic conditions and ionomycin treatment (related to Fig. 4). Bright field time lapse movie of progenitor cells cultured in isotonic condition (top-left), hypotonic media (top-right), hypotonic media supplemented with 1 μM ionomycin in suspension (bottom-left) or under 16 μm confinement (bottom-right). Hypotonic conditions supplemented with 1 μM ionomycin lead to rapid cell polarization and induce a stable-bleb cell transformation (bottom-left), but cells can migrate only under confined conditions (bottom-right).

**Movie S9.** The endoplasmic reticulum (ER) is immobilized under the nucleus in deformed cells under high confinement (related to Fig. 4). Time lapse TIRF movie of progenitor cells stained with ER tracker green under increasing mechanical confinement. At low mechanical confinement, the ER is mobile in the nucleus-plasma membrane contact area while for increasing confinement the ER is immobilized in the region underneath the nucleus. The yellow lines mark the nuclear area (visualized from bright field imaging).
